# Genome rearrangements and selection in multi-chromosome bacteria *Burkholderia spp*

**DOI:** 10.1101/319723

**Authors:** Olga O Bochkareva, Elena V Moroz, Iakov I Davydov, Mikhail S Gelfand

**Affiliations:** Kharkevich Institute for Information Transmission Problems, Moscow, Russia; Center for Data-Intensive Biomedicine and Biotechnology, Skolkovo Institute of Science and Technology, Moscow, Russia; Department of Ecology and Evolution & Department of Computational Biology, University of Lausanne, Lausanne, Switzerland; Swiss Institute of Bioinformatics, Lausanne, Switzerland; Faculty of Computer Science, Higher School of Economics, Moscow, Russia

**Keywords:** multi-chromosome bacteria, genome rearrangements, pan-genome, comparative genomics, strain phylogeny, positive selection

## Abstract

**Background:** The genus *Burkholderia* consists of species that occupy remarkably diverse ecological niches. Its best known members are important pathogens, *B. mallei* and *B. pseudomallei*, which cause glanders and melioidosis, respectively. *Burkholderia* genomes are unusual due to their multichromosomal organization.

**Results:** We performed integrated genomic analysis of 127 *Burkholderia* strains. The pan-genome is open with the saturation to be reached between 86,000 and 88,000 genes. The reconstructed rearrangements indicate a strong avoidance of intra-replichore inversions that is likely caused by selection against the transfer of large groups of genes between the leading and the lagging strands. Translocated genes also tend to retain their position in the leading or the lagging strand, and this selection is stronger for large syntenies. Integrated reconstruction of chromosome rearrangements in the context of strains phylogeny reveals parallel rearrangements that may indicate inversion-based phase variation and integration of new genomic islands. In particular, we detected parallel inversions in the second chromosomes of *B. pseudomallei* with breakpoints formed by genes encoding membrane components of multidrug resistance complex, that may be linked to a phase variation mechanism. Two genomic islands, spreading horizontally between chromosomes, were detected in the *B. cepacia* group.

**Conclusions:** This study demonstrates the power of integrated analysis of pan-genomes, chromosome rearrangements, and selection regimes. Non-random inversion patterns indicate selective pressure, inversions are particularly frequent in a recent pathogen *B. mallei*, and, together with periods of positive selection at other branches, may indicate adaptation to new niches. One such adaptation could be a possible phase variation mechanism in *B. pseudomallei*.

## Background

The genus *Burkholderia* comprises species from diverse ecological niches (Coenye and Vandamme, 2003). In particular, *B. mallei* and *B. pseudomallei* are pathogens causing glanders and melioidosis, respectively, in human and animals (Howe, Sampath, and Spotnitz, 1971); *B. glumae* is a pathogen of rice (Ham, Melanson, and Rush, 2011); *B. xenovorans* is an effective degrader of polychlorinated biphenyl, used for biodegradation of pollutants (Goris et al., 2004); *B. phytofirmans* is a plant-beneficial endophyte that may trigger disease resistance in the host plant (Frommel, Nowak, and Lazarovits, 1991). *Burkholderia* genomes are unusual due to their multichromosomal organization, generally comprised of two or three chromosomes.

By definition, the pan-genome of a genus or species is the set of all genes found in at least one strain (Tettelin et al., 2005). The core-genome is the set of genes shared by all strains; this gene set is usually used for accurate phylogenetic reconstruction. Genes that are not common for all considered strains but are not unique form the periphery part of a pan-genome. The pangenome of 56 *Burkholderia* genomes was estimated to exceed 40,000 genes with no sign of saturation upon addition of more strains, and the core-genome was approximately 1,000 genes (Ussery et al., 2009). A separate analysis of 37 complete *B. pseudomallei* genomes did not show saturation either (Spring-Pearson et al., 2015). The core-genome of *B. mallei* is smaller than that of *B. pseudomallei*, while the variable gene sets are larger (Losada et al., 2010).

In multi-chromosome bacterial species, the gene distribution among chromosomes is not random. The majority of genes necessary for the basic life processes usually are located in one (primary) chromosome. Other (secondary) chromosomes contain few essential genes and are mainly composed of niche-specific genes (Egan, Fogel, and Waldor, 2005). An exception is two circular chromosomes of *Rhodobacter sphaeroides* that share responsibilities for fundamental cell processes (Mackenzie et al., 2001). Genes from a secondary chromosome evolve faster than primary-chromosome genes and hence secondary chromosomes may serve as evolutionary test beds so that genes from secondary chromosomes provide conditional benefits in particular environments (Cooper et al., 2010). Secondary chromosomes usually evolve from plasmids (Egan, Fogel, and Waldor, 2005).

Several examples of gene translocations between chromosomes in *Burkholderia* are known, e.g., the translocation between the first and the third chromosomes in *B. cenocepacia* AU 1054, affecting many essential genes (Guo et al., 2010). Following interchromosomal translocation, genes change their expression level and substitution rate, dependent on the direction of the translocation (Morrow and Cooper, 2012).

Intra-chromosome genome rearrangements such as duplications, deletions, and inversions also play important roles in the bacterial evolution, as they strongly affect the chromosome organization and gene expression. Reconstruction of the history of genome rearrangements leads to a new class of phylogeny reconstruction algorithms (Alekseyev and Pevzner, 2009; Hu, Lin, and Tang, 2014). Chromosomal rearrangements often happen via recombination between repeated sequences, such as insertion (IS) elements (Raeside et al., 2014) and rRNA operons (Huang et al., 2008). Selection on inversion positions tends to preserve the size symmetry of the two replichores (regions of a circular chromosome between the origin and the terminus of replication), gene positions on the lagging/leading strand, and distances between genes and the origin of replication (Eisen et al., 2000; Kowalczuk et al., 2001). Sometimes inversions are mediated by inverted paralogs. Such inversions may lead to alternating expression of these paralogs; this mechanism is known as antigenic variation by which the organism may evade host immune responses (García-Pastor, Puerta-Fernández, and Casadesús, 2018).

Genome rearrangement played an important role in the *B. mallei* speciation. Genomic analyses of the first sequenced *B. pseudomallei* strains and their comparison with avirulent *B. thailandensis* have shown that both chromosomes are highly syntenic between the two species, with few large-scale inversions (Challacombe et al., 2014; Yu et al., 2006). In comparison to *B. pseudomallei*, *B. mallei* genomes harbor numerous IS elements that most likely have mediated the higher rate of rearrangements (Nierman et al., 2004). In particular, IS elements of the type IS407A had undergone a significant expansion in all sequenced *B. mallei* strains, accounting for 76% of all IS elements, and their chromosomes were dramatically and extensively rearranged by recombination across these elements (Losada et al., 2010). The genomic reduction of *B. mallei* following its divergence from *B. pseudomallei* likely resulted in its inability to live outside the host (Losada et al., 2010; Godoy et al., 2003).

Gene gains and losses also impact the pathogenicity of species and the adaptability of an organism. The loss of a type III secretion system (T3SS)-encoding fragment in *B. mallei* ATCC 23344, compared to *B. mallei* SAVP1, is responsible for the difference in the virulence between these strains (Schutzer et al., 2008). Another example is the loss of the L-arabinose assimilation operon by pathogens *B. mallei* and *B. pseudomallei* in comparison with an avirulent strain *B. thailandensis*. Introduction of the L-arabinose assimilation operon in a *B. pseudomallei* strain made it less virulent (Moore et al., 2004). Hence, although the mechanism is not clear, there may be a link between this operon and virulence. Acquisition of the atrazine degradation and nitrotoluene degradation pathways by *B. glumae* PG1, compared to *B. glumae* LMG 2196 and *B. glumae* BGR1, likely has resulted from an adaption since these toxic agents are used in the farming industry as a herbicide and a pesticide, respectively (Lee et al., 2016).

Here, we performed an integrated reconstruction of chromosome rearrangements for 127 complete *Burkholderia* strains, including inter-chromosome translocations, inversions, deletions/insertions, and single gene gain/loss events in the phylogenetic context. As the evolution of an obligate intracellular pathogen *B. mallei* from *B. pseudomallei* is a particular interest, we considered this branch of *Burkholderia* in additional detail, including analysis of pan-genome statistics and identification of genes evolving under positive selection.

## Methods

Available (as of 1 September 2016) complete genome sequences of 127 *Burkholderia* strains (Suppl. Table S1) were downloaded from the NCBI Genome database (Benson et al., 2013).

### Orthologs groups

We constructed orthologous groups using Proteinortho V5.13 with the default parameters (Lechner et al., 2011). To assign GO terms to genes, we used Interproscan (Jones et al., 2014). A GO term was assigned to an orthologous group, if it was assigned to at least 90% of genes in this group. To determine overrepresented functional categories, we used topGO v.3.6 package for R (Alexa and Rahnenfuhrer, 2016). Clusters of Orthologous Groups were predicted using eggNOG v4.5 database (Huerta-Cepas et al., 2016). Protein subcellular localization was predicted using PSORTb v3.02 web server (Yu et al., 2017). The expression level for each orthologous group was calculated based on RNA-seq data from (Lazar Adler et al., 2016). Using these data, we calculated RPKM, performed quantile normalization, and then calculated average value among samples.

### Pan-genome and core-genome size

To predict the number of genes in the *Burkholderia* pan-genome and core-genome, we used the binomial mixture model (Snipen, Almoy, and Ussery, 2009) and the Chao lower bound (Chao, 1987) implemented in the R-package Micropan (Snipen and Liland, 2015). To select the model better fitting the distribution of genes by the number of strains in which they are present, we used the Akaike information criterion with correction for a finite sample size (Akaike, 1974; Hurvich and Tsai, 1989).

### Phylogenetic trees

#### Trees based on nucleotide alignments

We performed codon alignment for each of the 2117 orthologous groups using Mafft version v7.123b (Katoh and Standley, 2013) and Guidance v2.01 (Penn et al., 2010). Four orthologous groups containing sequences scored below 0.8 were excluded from further analysis. Poorly aligned residues (guidance score below 0.8) were masked. The resulting sequences were concatenated and the tree was constructed with RAxML v8.2.9 (Stamatakis, 2014) using the GTR+Gamma model with 100 bootstrap runs. To ensure robustness of the tree construction we also performed calculations with 1000 replicates (see calculations at the GitHub repository).

#### Trees based on protein alignments

We used 1046 orthologous protein-coding genes from 127 genomes. We used Mafft v7.273 (Katoh and Standley, 2013) in the linsi mode to align genes belonging to one orthologous group. Concatenated protein-coding sequences were used to construct the tree. We used PhyML (Guindon et al., 2010) with the JTT model and discrete gamma with four categories and approximate Bayes branch supports.

#### Trees based on gene content

The gene content tree was constructed using the Neighbor Joining (NJ) algorithm based on the pairwise distance matrix 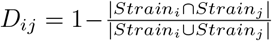, where *Strain_i_* is the set of orthologs belonging to a given strain *i*, ignoring paralogs.

#### Trees based on gene order

Trees based on gene order were built using the MLGO software (Maximum Likelihood for Gene-Order Analysis) with default parameters (Hu, Lin, and Tang, 2014).

#### Trees visualization

Phylogenetic trees were visualized with FigTree v1.4.2 (http://tree.bio.ed.ac.uk/software/figtree/) and *R* package Ape (Paradis, Claude, and Strimmer, 2004).

### Gene acquisition, loss, and translocations

We used GLOOME (Cohen et al., 2010) for the gain/loss analysis in the evolution non-stationary model with a variable gain/loss ratio. Other parameters were set based on character counts directly from the phyletic pattern.

To reconstruct gene translocations between chromosomes, we ordered universal single-copy orthologs and assigned a vector of ortholog presence to each strain. A component of this vector was the chromosome (1, 2, 3) harboring the ortholog in the strain. Then we subjected the obtained alignment of vectors to PAML 4.6 (Yang, 1997) for ancestral reconstruction with default parameters, except model = REV(GTR) and RateAncestor = 2.

### Synteny blocks and blocks rearrangements history

Synteny blocks for closely related strains were constructed using the Sibelia software (Minkin et al., 2013) with the minimal length of blocks being 5000 bp. We filtered out blocks observed in any single genome more than once. Synteny blocks for distant strains were constructed using the Drimm-Synteny program (Pham and Pevzner, 2010) based on locations of universal genes. The rearrangements histories for given trees topologies were constructed using the MGRA v2.2 server (Avdeyev et al., 2016).

To distinguish between inter- and intra-replichore inversions, the origins and terminators of replication for each chromosome of each strain were determined by the analysis of peaks in GC-skew plots combined with Ori-Finder predictions (Gao and Zhang, 2008). Statistical significance of over-representation of inter-replichore inversions was calculated as the probability of a given number of inter-replichore inversions in the set of inversions with the given lengths. The probability of occurrence of the origin or the terminator of replication within the inversion was calculated as the ratio of the inversion length to the replichore length.

### Detection of positive selection

We applied codon models for positive selection to orthologous groups common for the *B. mallei*, *B. pseudomallei*, *B. thailandensis*, *B. oklahomensis* clade. Given the low number of substitutions, it is usually not possible to reliably reconstruct a phylogenetic tree topology based on individual genes. On the other hand, given the high recombination rate, it is quite likely that gene evolutionary histories are slightly different between orthologous groups. To overcome these issues we first used statistical binning (Mirarab et al., 2014) to group genes with similar histories, and then applied a conservative approach to detect positive selection based on multiple tree topologies.

The procedure was implemented as follows. First, we constructed a phylogenetic tree for every gene using RAxML with the GTR+Gamma model and maximum likelihood with 100 bootstrap replicates. To ensure robustness of the tree construction we also performed calculations with 1000 replicates (see calculations at the GitHub repository). Genes with unexpectedly long branch lengths were filtered out (the maximum branch length > 0.1 or the sum of branch lengths > 0.3). Statistical binning was performed at the bootstrap incompatibility threshold of 95. For each of 25 obtained clusters we created a tree with bootstrap support using the concatenated sequence of orthologous groups belonging to the cluster.

We used two different methods to detect positive selection. The M8 vs M8a comparison allows for gene-wide identification of positive selection (Yang, 2007), while the branch-site model accounts for positive selection on a specific branch (Zhang, Nielsen, and Yang, 2005). Each test was performed six times using different trees: the maximum likelihood tree and five random bootstrap trees. We used the minimum value of the LRT (likelihood ratio test) statistic to avoid false identification of positive selection which could be caused by an incorrect tree topology.

For the branch-site model we tested each internal branch as a foreground branch one by one; we did not test terminal branches to avoid false positives caused by sequencing errors. The results of the branch-site tests were aggregated only in the case of bipartition compatibility. We considered only bipartitions that were present in at least three tests, we also computed the minimum value of the LRT statistic. The test results were mapped back to the species tree based on bipartition compatibility.

The strength of purifying selection is measured by *w*0 *<* 1 with smaller values corresponding to stronger purifying selection. The *w*2 > 1 parameter of the branch-site model captures positive selection, with higher values indicating stronger selection.

In both cases we used the chi-square distribution with one degree of freedom for the LRT to compute the *p*-value. Finally, we computed the *q*-value, all LRT values equal to zero were excluded from the test. We set the *q*-value threshold to 0.1.

### Statistical methods

To estimate dependencies between various parameters such as the expression level, localization in the first/second chromosome, localization on the leading/lagging strand, we used linear models (lm function, R v3.3.2). Additional parameters such as the sum of branch lengths, alignment length, and GC-content were included as they can affect the power of the method (Drummond et al., 2005). The parameters were transformed to have a bell-shaped distribution if possible: log(*x* + 1) for the expression levels, log(*x* + 10^*−*^6) for the LRT statistic, and log(*x*) for the alignment length, sum of branch lengths, standard deviation of GC-content, and *ω*0. Continuous variables were centered at zero and scaled so that the standard deviation was equal to one. This makes the linear model coefficients directly comparable. Outliers were identified in the residual plots and excluded from the model; the residual plots did not indicate abnormalities. For the linear models, we included potential confounding variables in the model, and kept only significant ones for the final linear model.

## Results and discussion

### Phylogeny and pan-genome analysis

The analysis of orthology for 127 *Burkholderia* strains yielded 757,526 orthologous groups containing two or more genes. 21,740 genes were observed in only one genome, some of them could result from misannotation. Alignments of 1024 single-copied common gene (hereinafter “core genes”) were used for construction of the phylogenetic tree (hereinafter “the basic tree”).

As the number of available *Burkholderia* genomes in GenBank is constantly increasing, we performed comprehensive pan-genome analysis. The pan-genome size for all strains is 48,000 genes with no signs of saturation, showing that the gene diversity of the *Burkholderia* species has not been captured yet (Fig. 1a). Based on these data, the binomial mixture model (Snipen, Almoy, and Ussery, 2009) predicts that, as more genomes are sequenced, the *Burkholderia* core-genome contains 457 genes, whereas the pan-genome size is 86,845. The number of new genes decreases with each new genome *n* at the rate *N* (*n*) = 2557*n*^−0,56^ confirming that the pan-genome is indeed open (Fig. S1a). Each new genome adds about 171 genes to the pan-genome. The Chao lower bound estimate (Chao, 1987)of the pan-genome size is 88,080. These results are consistent with the reported pan-genome size of 56 *Burkholderia* strains (Spring-Pearson et al., 2015). The core-genome size dependence on the number of analyzed strains is shown in Fig. 1b. The number of universal genes that are present in all strains saturates at about 1,050.

**Figure 1:**
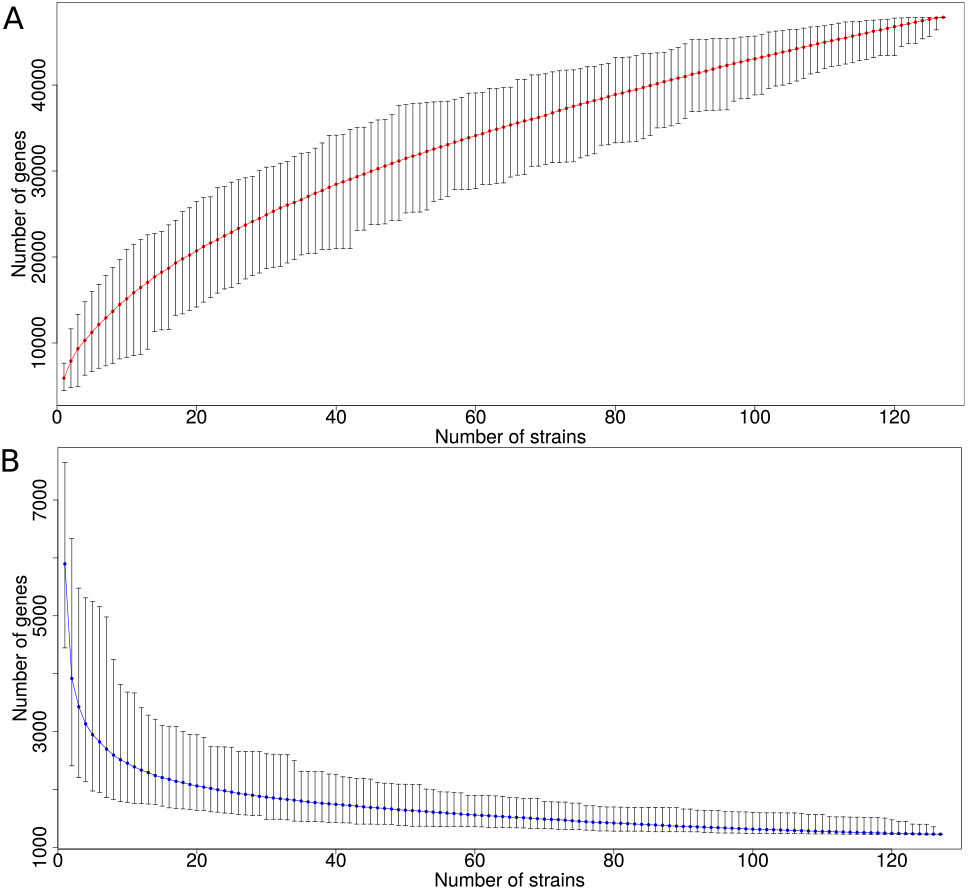
Number of genes as a function of the number of sequenced Burkholderia genomes. (A) The pan-genome size, that is, the number of all genes in sequenced strains. The number of new genes decreases with each new genome n at the rate N (n) = *2557*n^−^*^0,56^* confirming that the pan-genome is open. As the numbers of genomes n → ∞, the pan-genome size converges to 88,080 and the core-genome size converges to 457 genes. (B) The core-genome size, that is, the number of common genes in sequenced strains. The core-genome for a single strain (n = *1*) is defined as the number of genes in the strain.

Suppl. Fig. S2 and S3 show the core- and pan-genome size dependencies for *B. pseudomallei* and *B. mallei*, respectively. Their pan-genomes also have not reached saturation (*N* (*n*) = 788*n*^−0,53^ for *B. pseu-domallei* and *N* (*n*) = 867*n*^−0,87^ for *B. mallei*) (Suppl. Fig. S1b,c). These results are also consistent with the reported pan-genome size of 37 *B. pseudomallei* strains (Ussery et al., 2009).

The distribution of genes by the number of strains in which they are present has a typical U-shape form (Fig. 2), with numerous unique and universal genes and fewer periphery genes. We compared two models that are traditionally used for U-curve approximation, by the sum of three exponents (for unique genes, the periphery, and the universal genome, respectively) (Makarova et al., 2007) and by the sum of two power law functions, the first term describes the genes present in a few strains (almost unique), and the second term reflects the distribution of genes present in most strains (almost universal) (Gordienko, Kazanov, and Gelfand, 2013). Application the method of the least squares with the Akaike information criterion (AIC) revealed that the approximation by the sum of three exponents recapitulates the U-shape slightly better. This is consistent with the analysis of the *Streptococcus* pan-genome (Shelyakin et al., 2018), in which the sum of three exponents also has provided a better fit.

**Figure 2:**
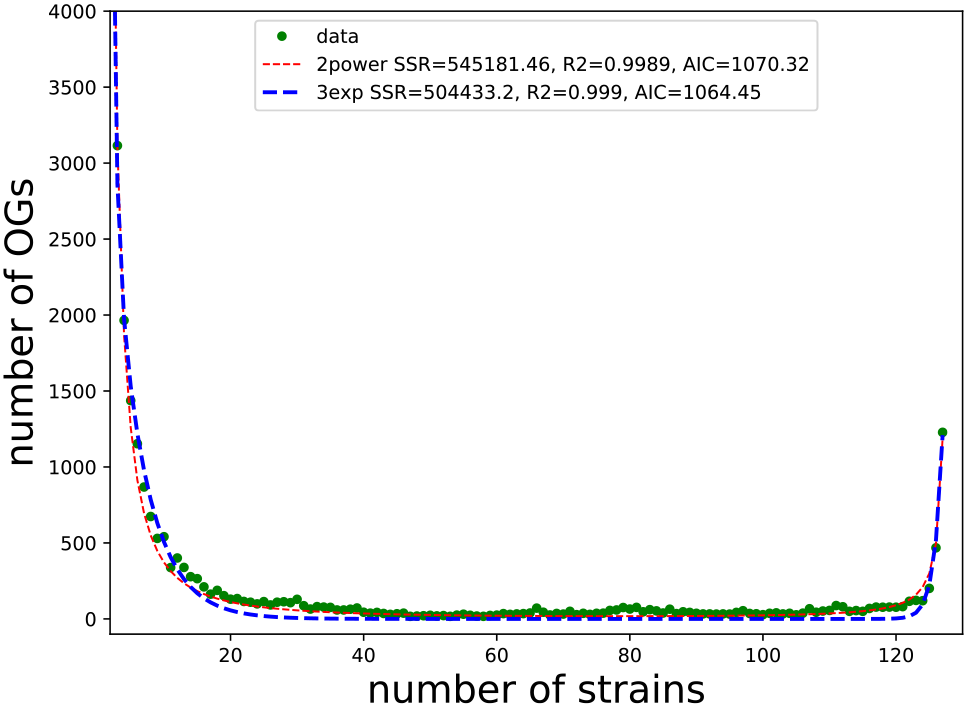
Distribution of ortholog groups (OGs) by the number of strains in which they are present. The blue line corresponds to the approximation by a sum of three exponents y *=* e^−*0.2*x *+8.4*^ *+* e^−*1.8*x*+11.6*^ *+* e^*0.85*x−*100.1*^; the red line corresponds to the approximation by a sum of two power functions y *= 21648.4*x^−^*^1.8^ + 1182.8(128* x)^−*1.2*^. Based on the Akaike information criterion (AIC), the approximation by the sum of three exponents recapitulates the U-shape slightly better.

### Genes acquisition and loss

Gains and losses of genes along the phylogenetic tree were assessed, excluding plasmid genes (Suppl. Fig. S4). *Burkholderia* species have experienced numerous gene gains and losses, that could explain their ecological diversity. In particular, a separate analysis of the *B. pseudomallei* group yielded considerable gene loss in the *B. mallei* clade. The genome reduction among the *B. mallei* strains is likely associated with the loss of genes redundant for obligate pathogens (Losada et al., 2010).

The basic tree and the gene content tree are largely consistent as the trees have the same clades with one major exception (Suppl. Fig. S5). In the gene content tree, *B. mallei* and *B. pseudomallei* form two distinct clusters, whereas in the basic tree monophyletic *B. mallei* are nested within paraphyletic *B. pseudomallei*. The former discrepancy could be due to the lifestyles of *B. mallei* and *B. pseudomallei*, as both species are pathogens of animals and possess specific sets of genes. Thus even if universal genes in some *pseudomallei* strains are closer to the orthologous genes in *mallei* than to genes in other *pseudomallei* strains, these species will be distant on the gene content tree due to species-specific genes.

Although the trees are comprised of the same clades, we observed numerous contradictions in strains positions. These contradictions are likely caused by clade-specific patterns of recombination and accessory gene exchange (Nandi et al., 2015).

### Rearrangements of universal single-copy genes

The gene distribution among the chromosomes (Table 1) for *Burkholderia* spp. is consistent with previous observations for other multi-chromosome bacteria (Egan, Fogel, and Waldor, 2005). At that, the majority of core genes belong to the first chromosome, ten-fold less core genes are in the second chromosome, and they are almost absent in the third chromosome, the only exception resulting from a large translocation from the first to the third chromosome in *B. cenocepacia* AU 1054 (Guo et al., 2010).

**Table 1:** Distribution of universal orthologs among the chromosomes. Each cell shows the number of universal genes out of the number of all genes in the chromosome. For species with more than one strain, the average number of genes and the standard deviation are shown.

| Species | The first chromosome | The second chromosome | The third chromosome |
| --- | --- | --- | --- |
| *B. mallei* | 930 out of 3135±103 | 116 out of 1842 ± 142 |  |
| *B. pseudomallei* | 954 out of 3498±151 | 92 out of 2536±178 |  |
| *B. thailandensis* | 954 out of 3626±216 | 92 out of 2562±109 |  |
| *B. oklahomensis* | 954 out of 3630±36 | 92 out of 2537±61 |  |
| *B. gladioli* | 964 out of 3924±146 | 82 out of 3026±100 |  |
| *B. glumae* | 964 out of 3385±120 | 82 out of 2524±365 |  |
| *B. vietnamiensis* | 961 out of 3055±139 | 85 out of 2411±420 |  |
| *B. cepacia* group | 962 out of 3205±146 | 84 out of 2528±413 | 0 out of 922±160 |
| *B. cenocepacia* AU 1054 | 747 out of 2965 | 84 out of 2472 | 215 out of 1040 |
| *B. cenocepacia* strain DDS 22E-1 | 961 out of 3296 | 84 out of 2831 | 1 out of 939 |
| *B. dolosa* AU0158 | 963 out of 3084 | 83 out of 1861 |  |
| *B. ubonensis* MCMB22 | 963 out of 3216 | 83 out of 3035 |  |
| *B. pyrrocinia* strain DSM 10685 | 963 out of 3157 | 84 out of 2714 | 0 out of 838 |
| *B. sp.* CCGE1001 | 968 out of 3545 | 78 out of 2420 |  |
| *B. sp.* CCGE1002 | 968 out of 3116 | 78 out of 2258 | 0 out of 1109 |
| *B. sp.* CCGE1003 | 967 out of 3463 | 79 out of 2525 |  |
| *B. sp.* HB1 | 967 out of 3481 | 79 out of 2743 |  |
| *B. sp.* KJ006 | 961 out of 2917 | 85 out of 2132 | 0 out of 930 |
| *B. sp.* OLGA172 | 967 out of 4023 | 79 out of 2998 |  |
| *B. sp.* PAMC 26561 | 964 out of 3034 | 82 out of 1437 |  |
| *B. sp.* PAMC 28687 | 960 out of 2991 | 83 out of 1367 | 3 out of 1509 |
| *B. sp.* RPE64 | 964 out of 2907 | 81 out of 1422 | 0 out of 853 |
| *B. sp.* RPE67 | 963 out of 2859 | 81 out of 1688 | 1 out of 1553 |
| *B. sp.* TSV202 | 954 out of 3645 | 92 out of 2536 |  |
| *B. sp.* YI23 | 963 out of 2769 | 81 out of 1539 | 1 out of 1364 |

Reconstruction of translocations of 1024 core genes between the chromosomes yielded 210 events (Fig. 3). The genomes of *B. cenocepacia* 895, *B. cepacia* strain LO6, and *B. contaminans* MS14 were not included in the rearrangement analysis due to likely artifacts of the genome assembly (See Suppl. Fig. S6). Thirty-eight events were reconstructed separately for *B. mallei* and *B. pseudomallei*.

**Figure 3:**
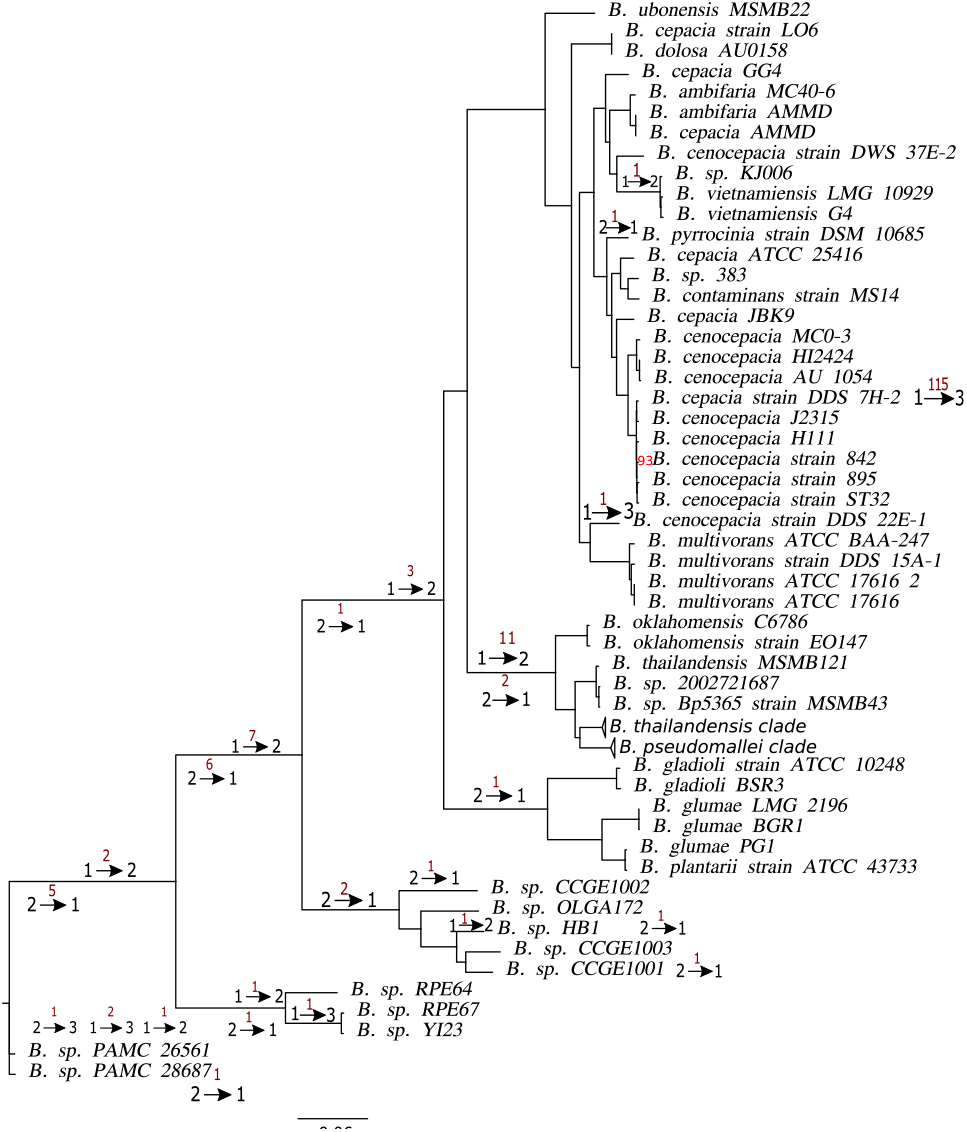
Translocations in Burkholderia spp. The phylogenetic tree of Burkholderia is constructed based on the protein sequence similarity of single-copy universal genes. The bootstrap support is shown for branches where it is < *100*. The red numbers above the arrows show the number of genes translocated between chromosomes on the tree branches; the black numbers mark chromosomes that have been involved in the transfer, the arrows show the direction of the transfer.

There was no statistically significant overrepresentation of GO categories in the set of translocated genes. Six genes have been translocated independently on different tree branches twice or more times, encoding aldo/keto reductase (IPR020471), HTH-type transcriptional regulator *argP* (IPR017685), gamma-glutamyltranspeptidase (IPR000101), acid phosphatase *acpA* (IPR017768), tryptophan synthase beta subunit-like PLP-dependent enzyme (IPR036052), *tonB*-dependent receptor. The reconstructed common ancestor of *Burkholderia* has 965 universal single-copy genes in the first chromosome, and 81, in the second chromosome.

We analyzed intra-chromosomal rearrangements that involve the core genes using only one representative strain from clades with closely related species. The core genes were grouped into 87 synteny blocks that contained two or more core genes in the same order in all analyzed genomes. The rearrangements history yielded no parallel events except parallel translocations between chromosomes described above. While one could expect that changes in the lifestyle and population bottlenecks could increase mutation and recombination rates simultaneously, no correlation between the number of rearrangements and the average mutation rates of the core genes was observed (data not shown).

We then reconstructed the detailed history of rearrangements in specific clades. A large number of available genomes of closely related bacterial strains allows one to consider micro-rearrangements in the evolutionary context, revealing parallel events that may indicate the action of antigenic variation (Shelyakin et al., 2018). An integrated analysis of sequence-based and inversion-based trees enhances the resolution of the phylogenetic reconstruction in the case of a high rate of genome rearrangements in a population (Bochkareva et al., 2018).

### Rearrangements in the *B. cepacia* group

For 27 strains of the *cepacia* group, the average coverage of chromosomes by synteny blocks was 50% for the first, 30% for the second, and less than 10% for the third chromosome. This agrees with the preferred location of universal genes discussed above. Hereafter, the third chromosomes are not considered due to their low conservation. Fixing the tree to the basic one, we reconstructed 17 inversions and 574 insertion/deletion events. The topology of the phylogenetic tree based on the order of synteny blocks (Suppl. Fig. S7c) is not consistent with the basic tree and a majority of deep nodes have low bootstrap support that may be explained by numerous parallel gain/loss events.

Only one parallel inversion of length 530 kb was found in the first chromosome of *B. cenocepacia* AU 1054 and *B. cenocepacia* J2315, the inversion break-points formed by the 16S-23S rRNA locus. In order to distinguish between truly parallel events and homologous recombination between these strains, we constructed a tree based on proteins encoded by genes from the inverted fragment. *B. cenocepacia* AU 1054 and *B. cenocepacia* J2315 did not change their position in the tree, and, in particular, did not cluster together (data not shown). Hence, this block was not subject to homologous recombination between these strains.

Two non-universal synteny blocks were found in different chromosomes in different strains. One block with length 8.5 kb is located in the first chromosome of *B. cenocepacia* MC0-3 and in the second chromosome of *B. cepacia* ATCC 25416. This genomic island contains five genes that belong to the iron uptake pathway, and an AraC family protein. Some genes of this cassette were also found in other *Burkholderia* species (Fig. 4a).

**Figure 4:**
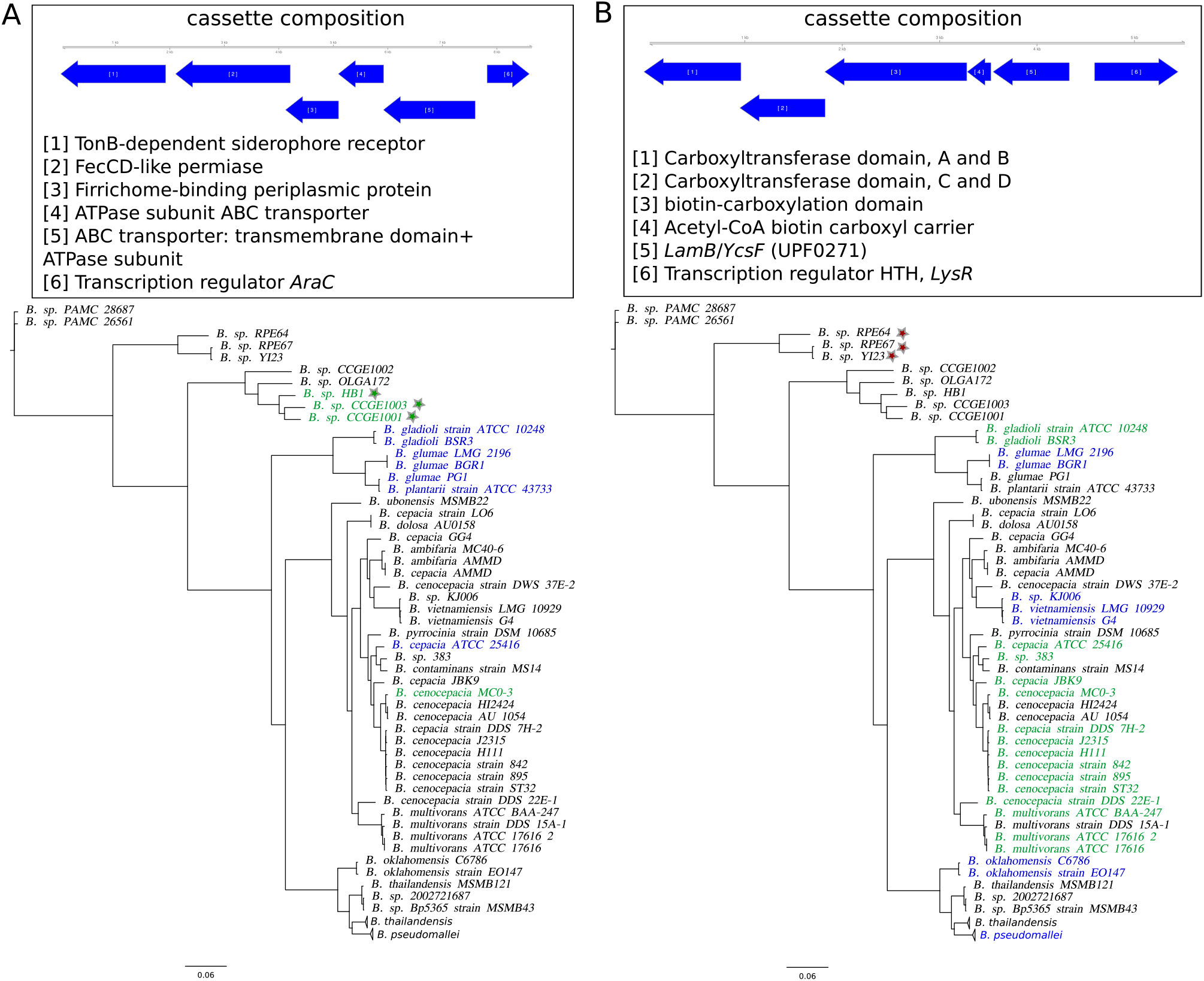
Phyletic patterns of two genomic islands detected in the B. cepacia group. Strains with the genomic island in the first chromosome are marked by green; strains with the genomic island in the second chromosome are marked by blue; red indicates location on plasmids. Strains with an incomplete cassette are marked by stars. Phyletic patterns are shown on the basic tree.

Another block with length 5.5 kb was found only in 17 of 30 strains belonging to the *cepacia* group (Fig. 4b). This island contains four genes forming the acetyl-CoA carboxylase complex, glycoside hydrolase (GO:0005975 carbohydrate metabolic process), and a LysR family protein. The island is found in all *B. mallei*, *B. pseudomallei*, *B. oklahomensis*, *B. glumae*, *B. gladioli* and is absent in *B. thailandensis* and other strains. Its presence in different chromosomes and differences between the tree of this cassette (Suppl. Fig. S8) and the basic tree indicate that this genomic island is spreading horizontally.

### Rearrangements in the *B. mallei* clade

For fifteen *B. mallei* strains and two *B. pseudomallei* used as outgroups, we constructed 104 common synteny blocks in both chromosomes. Only one block with length 40 kb, comprising 24 universal genes, was translocated in the *B. mallei* clade. This block is bounded by IS elements and rRNAs that may indicate that this translocation resulted from recombination between chromosomes.

This indicates that in these strains translocations between chromosomes are rare in comparison to within-chromosome rearrangements. Fixing the tree to the basic one, we reconstructed 88 inversions in the first chromosomes and 27 inversions in the second ones (Fig. 5). The reconstruction yields nine parallel events in the first chromosomes and three, in the second ones. The boundaries of the inversions are formed by repeated sequences (transposases).

**Figure 5:**
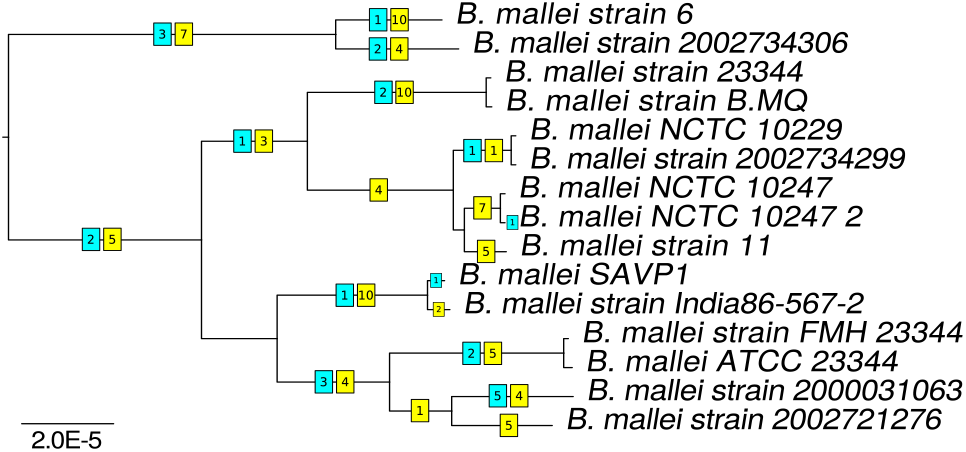
Inversions in the B. mallei clade. The numbers of inversions on branches are shown in squares on the basic tree. Yellow and blue color marks inversions in the first and second chromosomes, respectively.

**Figure 6:**
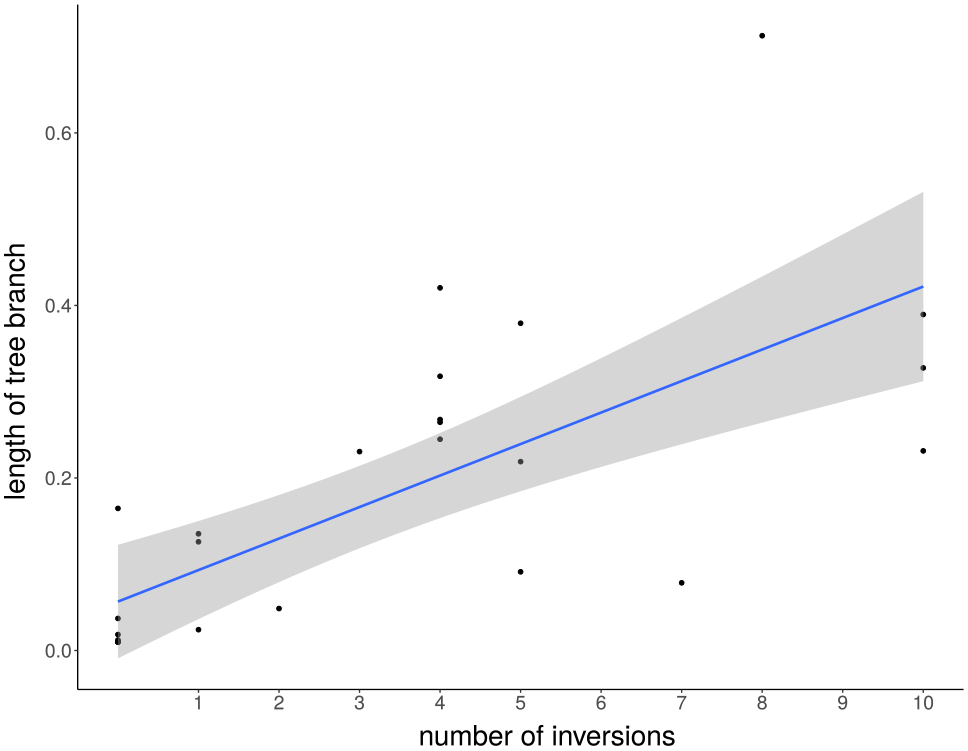
The rearrangements rate as a function of the mutation rate for B. mallei. Each dot corresponds to a branch in the basic phylogenetic tree (Fig. 5).

To test the possibility the contradictions between the tree topology and the inversion history had been caused by homologous recombination, we constructed trees based on genes involved in these events. For all inverted sequences, strains do not change their positions in the tree (data not shown). Therefore, we suppose that parallel events were caused by active intragenome recombination linked to a limited number of repeated elements.

We applied maximum likelihood optimization methods to obtain a topology based on the universal gene order. The optimized topology (Suppl. Fig. S7a) yielded

### Rearrangements in *B. pseudomallei*

The gene order in 51 strains of *B. pseudomallei* turned out to be significantly more stable than that in *B. mallei*, as only three inversions were reconstructed in the first chromosomes, and five, in the second chromosomes (Fig. 7a). Moreover, the average coverage of chromosomes by synteny blocks was more than 90% for the first and 80% for the second chromosomes, revealing a stable order and gene content. Two blocks with length about 20-25 kb are swapped in *B. pseudomallei* K42 that is likely to be an assembly artifact.

**Figure 7:**
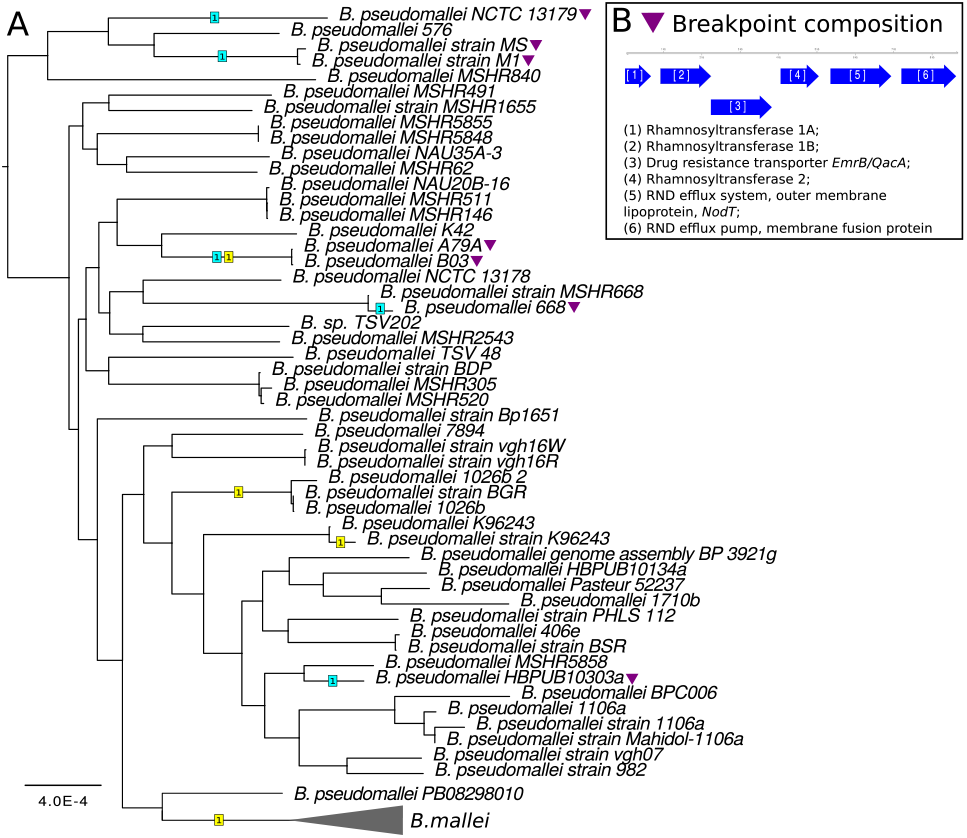
Inversions in B. pseudomallei. (A) The numbers of inversions on branches are shown in squares on the basic tree. Yellow color corresponds to inversions in the first chromosomes, blue color corresponds to the second chromosomes. The parallel inversion is marked by color triangles. (B) The breakpoint composition of the parallel inversion in the second chromosomes in B. pseudomallei.

Inversions in the second chromosomes with length about 1.3 Mb have the same boundaries for all seven strains despite the fact that they are located at distant branches of the phylogenetic tree (Fig. 7b). Break-points of these inversions are formed by six genes encoding (1,2) rhamnosyltransferase type 1 A,B; (3) drug resistance transporter (*mrB/QacA* subfamily); (4) rhamnosyltransferase type II; (5,6) components of a RND efflux system, outer membrane lipoproteins *nodT* and *emrA*.

The observed parallel inversion between paralogous genes encoding surface antigen protein might indicate the action of an antigen variation mechanisms leading to phenotype diversification. While direct confirmation of this mechanism requires analysis of transcripts, parallel, independent inversions are a good lead for subsequent experimental validation (Shelyakin et al., 2018).

### Rearrangements in the *B. thailandensis* clade

For 15 strains *B. thailandensis*, we constructed 56 synteny blocks in both chromosomes. Two strains of *B. oklahomensis* and one *B. pseudomallei* were used as out-groups. The average coverage by blocks was 75% for the first, and 50% for the second chromosomes. Fixing the tree topology to the basic tree, we reconstructed 18 inversions and 265 insertion/deletion events (Fig. 8). *B. thailandensis* has a higher rate of inversions and deletions than *B. oklahomensis* and *B. pseudomallei*. The reconstruction yields two parallel events in the first chromosomes and one, in the second ones. The boundaries of these inversions are formed by repeated sequences (transposases). For all inverted sequences, strains do not change their position in the trees based on sequences similarities of genes involved in these events (data not shown).

**Figure 8:**
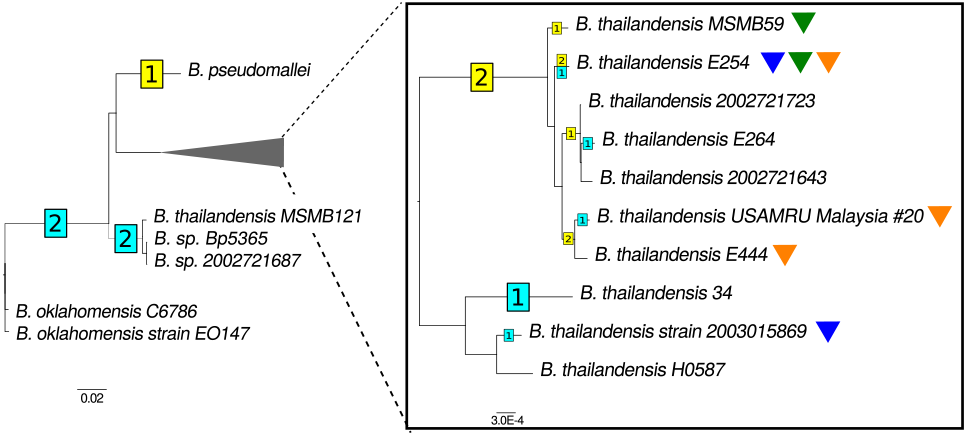
Inversions in B.thailandensis. The number of inv sions on branches are shown in squares on the ba tree. The area indicated by the gray triangle at the left panel is zoomed in the right panel. Yellow and blue colors mark inversions in the first and second chromosomes, respectively. Parallel inversions are marked by colored triangles.

The topology of the phylogenetic tree based on the order of synteny blocks (Suppl. Fig. S7b) is largely consistent with the basic tree, the only exception being a changed position of *B. thailandensis* E254 caused by parallel inversions.

Two non-universal, non-trivial translocated synteny blocks were found. One is a block with length 38 kb in the first chromosome in *B. pseudomallei*, the second chromosome in *B. oklahomensis*, and absent in the *B. thailandensis* genomes. This block is comprised of genes linked with amino acids metabolism. The second block is a parallel phage insertion with length 9 kb in the first chromosome of *B. oklahomensis* strain EO147 and in the second chromosome of *B. thailandensis* 2003015869.

### Selection regimes

As the evolution of species from the *B. thailandensis*, *B. pseudomallei*, and *B. mallei* clade is of particular interest due to dramatic changes in their lifestyle, including an adaptation to intra-cellular one, for these strains we identified genes evolving under positive selection. 1842 single-copy genes common for the *B. oklahomensis, B. thailandensis, B. pseudomallei, B. mallei* clade were tested. We detected 197 genes evolving under positive selection using the M8 model (Suppl. Table S2). No GO categories were significantly overrepresented but we observed overrepresentation of outer membrane proteins (permutation test, *p*-value=0.03) consistent with observations in other bacterial species (Cao et al., 2017; Xu, Chen, and Zhou, 2011).

To identify branch-specific positive selection, we used the branch-site test. In total, we identified seventeen events (Table 2), twelve of which we successfully mapped to the basic tree (Fig. 9). In the remaining five cases (flagellar hook protein FlgE, porin related exported protein, penicillin-binding protein, phosphoenolpyruvate-protein kinase and cytidylate kinase), the detected branches (bipartitions) of the gene trees were incompatible with the basic tree, and thus could not be mapped to it.

**Table 2:**
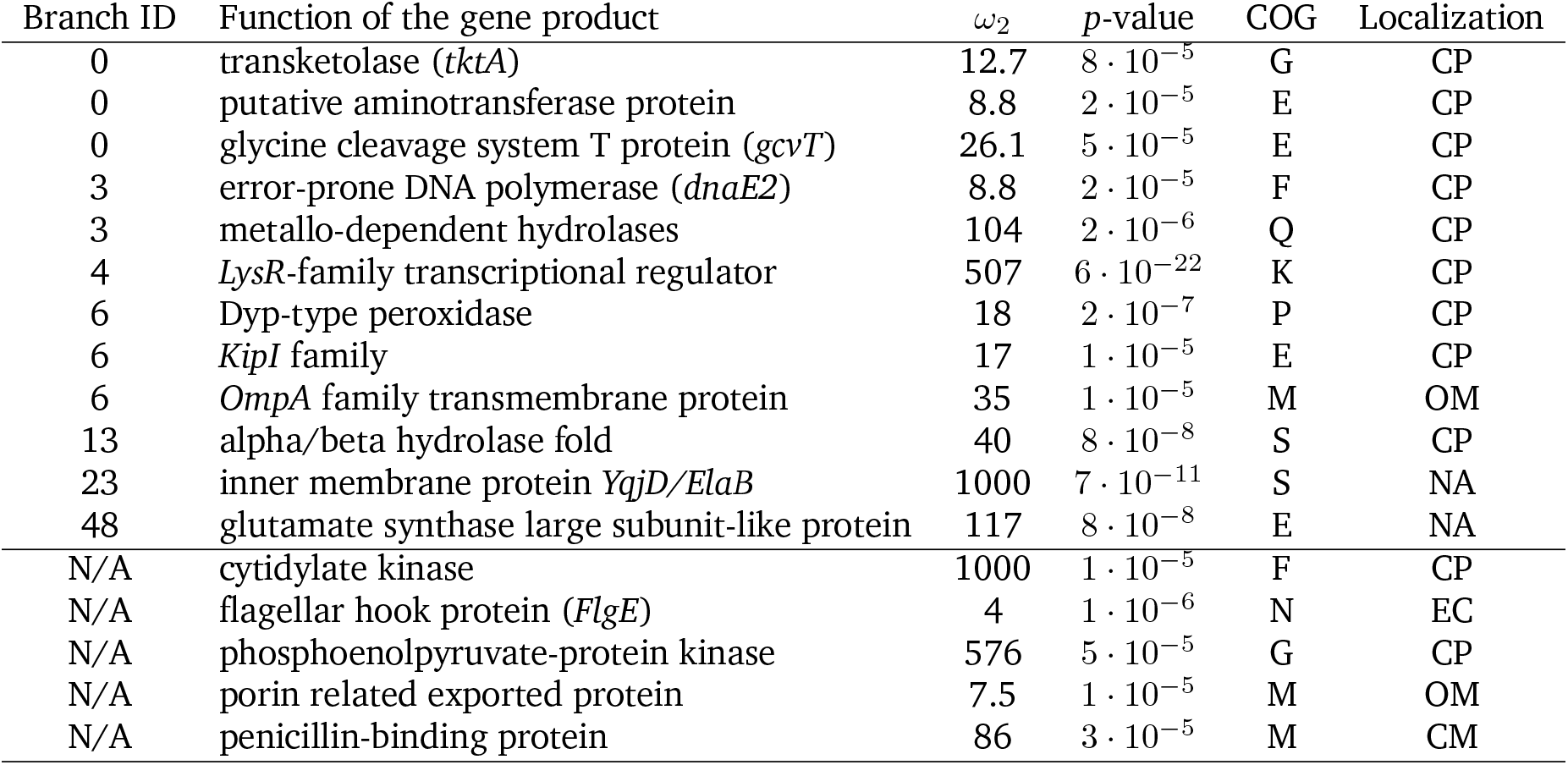
Genes evolving under branch-specific positive selection. The COG categories are coded as follows: K, transcription; M, cell wall/membrane biogenesis; N, Cell motility; G, carbohydrate transport and metabolism; E, amino acid transport and metabolism; F, nucleotide transport and metabolism; P, inorganic ion transport and metabolism; Q, secondary metabolites biosynthesis, transport and catabolism; R General function prediction only. The localization is coded as follows: CP, Cytoplasmic; OM, outer membrane; EC, Extracellular; CM, CytoplasmicMembrane; NA, unknown (these proteins may have multiple localization sites).

| Branch ID | Function of the gene product | $\omega_2$ | $p$-value | COG | Localization |
| --- | --- | --- | --- | --- | --- |
| 0 | transketolase (*tktA*) | 12.7 | $8 \cdot 10^{-5}$ | G | CP |
| 0 | putative aminotransferase protein | 8.8 | $2 \cdot 10^{-5}$ | E | CP |
| 0 | glycine cleavage system T protein (*gcvT*) | 26.1 | $5 \cdot 10^{-5}$ | E | CP |
| 3 | error-prone DNA polymerase (*dnaE2*) | 8.8 | $2 \cdot 10^{-5}$ | F | CP |
| 3 | metallo-dependent hydrolases | 104 | $2 \cdot 10^{-6}$ | Q | CP |
| 4 | *LysR*-family transcriptional regulator | 507 | $6 \cdot 10^{-22}$ | K | CP |
| 6 | Dyp-type peroxidase | 18 | $2 \cdot 10^{-7}$ | P | CP |
| 6 | *KipI* family | 17 | $1 \cdot 10^{-5}$ | E | CP |
| 6 | *OmpA* family transmembrane protein | 35 | $1 \cdot 10^{-5}$ | M | OM |
| 13 | alpha/beta hydrolase fold | 40 | $8 \cdot 10^{-8}$ | S | CP |
| 23 | inner membrane protein *YqjD/ElaB* | 1000 | $7 \cdot 10^{-11}$ | S | NA |
| 48 | glutamate synthase large subunit-like protein | 117 | $8 \cdot 10^{-8}$ | E | NA |
| N/A | cytidylate kinase | 1000 | $1 \cdot 10^{-5}$ | F | CP |
| N/A | flagellar hook protein (*FlgE*) | 4 | $1 \cdot 10^{-6}$ | N | EC |
| N/A | phosphoenolpyruvate-protein kinase | 576 | $5 \cdot 10^{-5}$ | G | CP |
| N/A | porin related exported protein | 7.5 | $1 \cdot 10^{-5}$ | M | OM |
| N/A | penicillin-binding protein | 86 | $3 \cdot 10^{-5}$ | M | CM |

**Figure 9:**
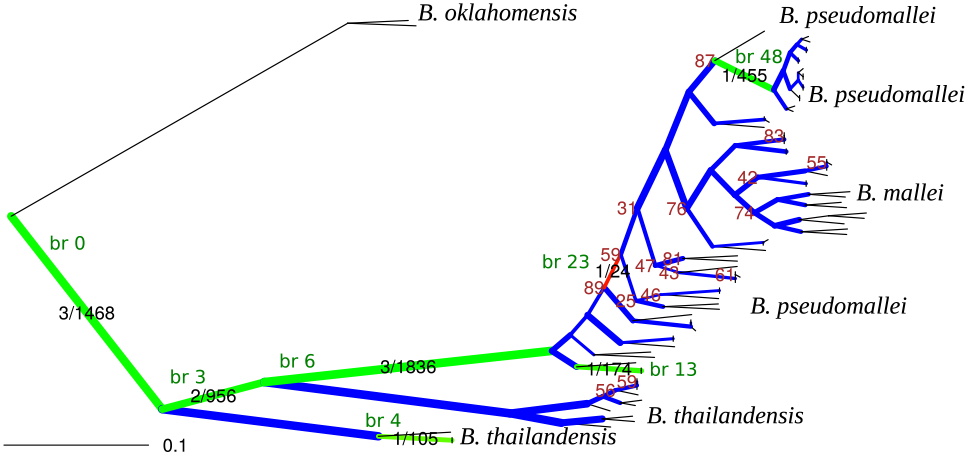
Phylogenetic species tree with detected events of positive selection. The phylogenetic tree is constructed based on the nucleotide sequence similarity of single-copy universal genes. The branch lengths are transformed using the square root. The bootstrap support is shown only for branches where it is < *90*. The branch thickness reflects the number of positive selection tests mapped on this branch. Color indicates the fraction of significant tests (blue=0; green, low rate; red, high rate), this number and the total number of tests are indicated on branches were positive selection has been detected. Branches with detected episodes of positive selection are marked by IDs in green squares that correspond to branches ID in Table 2. The tree with full strain names is shown in Suppl. Fig. S9.

Outer membrane proteins such as the flagellar hook protein FlgE, porin-related exported protein, OmpA family protein can serve as targets for the immune response. Moreover, OmpA is known to be associated with virulence, being involved in the adhesion and invasion of host cells, induction of cell death, serum and antimicrobial resistance, and immune evasion (Sousa et al., 2012). Error-prone DNA polymerase has a lower replication accuracy, and, thus, a higher mutation rate. Positive selection on this polymerase might be a result of adaptation to a new life style. Bacterial transcription factors are known to enable rapid adaptation to environmental conditions, that might explain strong positive selection of the LysR-family transcriptional regulator.

The majority of genes evolving under positive selection have been identified in the longest branches; accordingly, the fraction of events is higher in these branches. This might indicate rapid adaptation to new ecological niches during species formation. However, the branch-site test for positive selection is more powerful on longer branches, and the position of a branch in the tree might affect the power (Yang and Reis, 2011). Hence, overrepresentation of positive selection events can be related to the power of the method, and does not necessary indicate the higher number of genes affected by positive selection on these branches.

We used linear modeling to identify determinants affecting purifying selection (Table 3). The strongest observed correlation is that highly expressed genes tend to evolve under stronger purifying selection, which is also consistent with previous observations (Cooper et al., 2010). The expression levels in our dataset are higher for the first chromosome (Table S3), which is consistent with observations for other multi-chromosome bacterial species (Dryselius et al., 2008). While the effect of correlation is not particularly high, the correlation is strongly statistically significant. This indicates that despite high stochasticity of mRNA expression, there is a statistically strong association between the expression level and gene localization and average GC content.

**Table 3:** The linear model of average ω (negative selection, estimated using M8), non-significant variables removed from the model. For the full model see Suppl. Table 3. The model p-value is < *2.2* • *10*^−*16*^; *the adjusted R^2^* is 0.2882.

| | Estimate | Std. Error | $t$-value | $p$-value |
| --- | --- | --- | --- | --- |
| Alignment length | -0.085 | 0.025 | -3.397 | 0.000704 |
| Average expression level | -0.079 | 0.026 | -3.023 | 0.002561 |
| Sum of branch lengths | 0.518 | 0.026 | 19.638 | $< 2 \cdot 10^{-16}$ |

Longer genes tend to experience stronger purifying selection that is consistent with previously shown negative correlation between the *d_N_ /d_S_* value and the median length of protein-coding genes in a variety of species (Novichkov et al., 2009). However, this observation also could be explained by the greater power in detecting strong negative selection in longer genes, similarly to the increase in the power when detecting positive selection for longer genes (Yang and Reis, 2011).

### Selection on recombination events

In many bacteria, within-replichore inversions, that is, inversions with endpoints in the same replichore, have been shown to be relatively rare and significantly shorter than inter-replichore inversions (Darling, Miklós, and Ragan, 2008; Repar and Warnecke, 2017). The pattern of inversions reconstructed for both chromosomes in *B. mallei* is consistent with both of these observations.

Inter-replichore inversions are overrepresented in the first (*p*-value *<* 10^−33^) and the second (*p*-value *<* 10^−30^) chromosomes. The lengths of inter-replichore inversions have a wide distribution up to the full replichore size (Fig. 10a), whereas the observed within-replichore inversions mainly do not exceed 15% of the replichore length. We observed only two longer inversions, both in *B. mallei* FMH23344. These inversions overlap with each other and may be explained by a single translocation event. This strong avoidance of inter-replichore inversions is probably caused by selection against gene movement between the leading and the lagging strands (Zhang and Gao, 2017).

**Figure 10:**
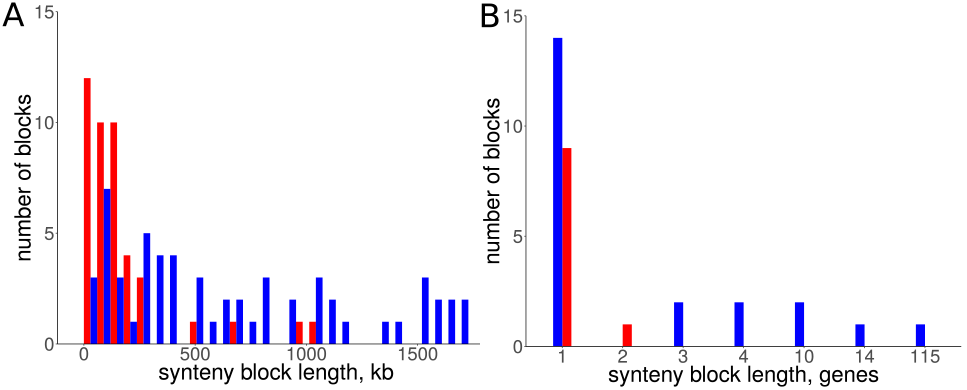
Histograms of lenghts of rearranged synteny blocks. (A) Blocks inverted in the first chromosomes in B. mallei; (B) blocks translocated between chromosomes in Burkholderia spp. Blue color corresponds to synteny blocks that have retained their position with respect to the leading/lagging strand; red color corresponds to synteny blocks that changed the strand.

The reconstruction of translocations also revealed that genes tend to retain their position on the leading or lagging strand (two-sided Binomial test, *p*-value=0.03, Fig. 10b). Moreover, all blocks of more than three genes retain their positions. We have not observed any difference in the level of purifying selection between genes translocated from the leading and lagging strands.

### Conclusions

The rearrangement rates differ dramatically in the *Burkholderia* species; from a couple of inversions in *B. pseudomallei* strains to dozens of events in *B. mallei* strains. Young pathogens such as *Yersinia pestis*, *Shigella* spp., *B. mallei* are known to have a particularly high rates and a variety of mobile elements that may be explained by fast evolution under changed selection pressure in new conditions, bottlenecks in the population history, and weaker selection against repetitive elements due to the decreased effective population size (Mira, Pushker, and Rodriguez-Valera, 2006). Accumulation of IS elements is most likely responsible for frequent genome rearrangement and strong genome reduction in *B. mallei* (Nierman et al., 2004). The tree based on the alignment of universal genes and the gene content tree also show some differences, caused by excessive gene gains and losses at some branches, most notably, gene loss in *B. mallei* following a drastic change of the lifestyle.

Integrated reconstruction of chromosome rearrangements in the context of strains phylogeny reveals parallel rearrangements. In particular, we detected parallel inversions in the second chromosomes of *B. pseudomallei* with breakpoints formed by genes encoding membrane components of multidrug resistance complex, that may be linked to a phase variation mechanism. Two genomic islands, spreading horizontally between chromosomes, were detected in the *B. cepacia* group. Hence, evolutionary and functional analysis of parallel rearrangements identifies possible cases of phase variation by inversions and integration of new genomic islands that is especially important for the micro-evolution of pathogens.

The observed strong avoidance of large intra-replichore inversions is likely caused by selection against transfer of large groups of genes between the leading and the lagging strands. At that, translocated genes also tend to retain their position in the leading or the lagging strand and this selection is stronger for large syntenies. This result is consistent with the inversion pattern in other bacterial species (Darling, Miklós, and Ragan, 2008) that may be explained, in particular, by over-presentation of highly expressed genes on the leading strand (Price and Arkin, 2005).

Overall, this study demonstrates the strength of integration of diverse approaches to the analysis of bacterial genomic evolution.

## Declarations

### Availability of data and materials

The datasets supporting the conclusions of this article and used *ad hoc* scripts are available via the link https://github.com/OlgaBochkaryova/burkholderia-genomics.

## Author’s contributions

MSG conceived the study, OOB, EVM and MSG designed the study; EVM, OOB and IID developed the methods, analyzed the data; EVM, OOB and IID wrote the manuscript, MSG reviewed the paper. All authors read and approved the final version of the manuscript.

## Ethics approval and consent to participate

Not applicable

## Consent for publication

Not applicable

## Acknowledgements

We thank Pavel Shelyakin for valuable comments. Analysis of parallel inversions was performed by Alisa Rodionova at the Summer School of Molecular and Theoretical Biology (Barcelona, 2016), supported by the Zimin Foundation.

## Additional Files

Additional file Fig. S1 — The number of new genes added to the pangenome upon addition of new strains. (a) *Burkholderia* spp., (b) *B. pseudomallei*, and (c) *B. mallei*. The number of specific genes is plotted as a function of the number (n) of strains sequentially added (see (Donati et al., 2010)). For each n, points are the values obtained for the different strain combinations; red symbols are the average of these values. The superimposed line is a fit with a decaying power law *y = A * n^B^*

Additional file Fig. S2 — Pan-genome (a) and coregenome (b) size of *B. pseudomallei* strains.

Additional file Fig. S3 — Pan-genome (a) and coregenome (b) size of *B. mallei* strains.

Additional file Fig. S4 — Gene flow during *Burkholderia* evolution. Red and blue numbers are, respectively, the numbers of gained and lost genes on a given branch.

Additional file Fig. S5 — Comparison the topologies of phylogenetic trees based on the protein sequence similarity of single-copy universal genes and the gene content.

Additional file Fig. S6 — Whole-genome alignments of *cepacia* strains that were not included in the rearrangement analysis due to likely artifacts of the genome assembly. (a) *Burkholderia* sp. 383 and *B. cepacia* strain LO6 (b) *Burkholderia* sp. 383 and *B. contaminans* strain MS14, (c) *Burkholderia* sp. 383 and *B. cenocepacia* strain 895, (d) *B. cepacia* strain LO6 and *B. cenocepacia* strain 895.

Additional file Fig. S7— Tanglegrams showing the differences between the tree topologies based on the protein sequence similarity of single-copy universal genes and the tree topologies based on the synteny blocks arrangements. (a) *B. mallei* clade; (b) *B. thailandensis* clade; (c) *B. cepacia* group.

Additional file Fig. S8— Phylogenetic tree constructed based on the concatenation of alignments of genes forming the genomic island.

Additional file Fig. S9— Phylogenetic tree showing detected events of positive selection.

Additional file Table S1 — Analyzed *Burkholderia* strains with genomes characteristics.

Additional file Table S2 — Genes evolving under positive selection.

Additional file Table S3 — Linear models (a) of average *ω* (negative selection, estimated using M8); (b) expression level (Lazar Adler et al., 2016).

